# DRP1-mediated mitochondrial fission is essential to maintain cristae morphology and bioenergetics

**DOI:** 10.1101/2021.12.31.474637

**Authors:** Gabriella L. Robertson, Stellan Riffle, Mira Patel, Andrea Marshall, Heather Beasley, Edgar Garza Lopez, Jianqiang Shao, Zer Vue, Maria S. Stoll, Sholto de Wet, Rensu P. Theart, Ben Loos, Antentor Hinton, Jason A. Mears, Vivian Gama

## Abstract

Mitochondria and peroxisomes are both dynamic signaling organelles that constantly undergo fission. While mitochondrial fission is known to coordinate cellular metabolism, proliferation, and apoptosis, the physiological relevance of peroxisome dynamics and the implications for cell fate are not fully understood. DRP1 (dynamin-related protein 1) is an essential GTPase that executes both mitochondrial and peroxisomal fission. Patients with *de novo* heterozygous missense mutations in the gene that encodes DRP1, *DNM1L (Dynamin 1 Like)*, present with encephalopathy due to defective mitochondrial and peroxisomal fission (EMPF1). EMPF1 is a devastating neurodevelopmental disease with no effective treatment. To interrogate the mechanisms by which DRP1 mutations cause cellular dysfunction, we used human-derived fibroblasts from patients with mutations in DRP1 who present with EMPF1. As expected, patient cells display elongated mitochondrial morphology and lack of fission. Patient cells display a lower coupling efficiency of the electron transport chain, increased proton leak, and upregulation of glycolysis. In addition to these metabolic abnormalities, mitochondrial hyperfusion results in aberrant cristae structure and hyperpolarized mitochondrial membrane potential, both of which are tightly linked to the changes in metabolism. Peroxisome structure is also severely elongated in patient cells and results in a potential functional compensation of fatty acid oxidation. Understanding the mechanism by which DRP1 mutations cause these metabolic changes will give insight into the role of mitochondrial dynamics in cristae maintenance and the metabolic capacity of the cell, as well as the disease mechanism underlying EMPF1.

## Introduction

Mitochondrial dynamics are controlled and executed by a vast group of proteins and other organelles. The underlying morphology of the mitochondria is an important determinant for the ability of this organelle to regulate a variety of cellular processes, such as apoptosis, cell fate, lipid biosynthesis, and epigenetic modifications (Bayraktar et al., 2019; Iwata et al., 2020; Khacho et al., 2016; King et al., 2021; Prigione and Adjaye, 2011; Rasmussen et al., 2020; Vantaggiato et al., 2019). Mitochondrial fusion is primarily catalyzed by the large GTPases Mitofusin 1 and Mitofusin 2 at the outer mitochondrial membrane (Chen et al., 2003). The GTPase OPA1 (Optic Atrophy 1) is responsible for executing fusion of the inner mitochondrial membrane (Alexander et al., 2000; Delettre et al., 2000). Mitochondrial fission, mediated through the large GTPase DRP1 (Dynamin-related protein 1), is orchestrated by a complex sequence of events (Fonseca et al., 2019; Smirnova et al., 2001). The first steps of mitochondrial fission include contact with the endoplasmic reticulum (ER) and replication of the mitochondrial genome (Friedman et al., 2011; Lewis et al., 2016). This is followed by contact and pre-constriction by actin filaments, which are associated with the ER membrane (Chakrabarti et al., 2018; Hatch et al., 2016, 2014; Ji et al., 2017, 2015; Li et al., 2015). Fission is also regulated through adaptor proteins at the outer mitochondrial membrane, including MiD49 (Mitochondrial Dynamics Protein 49), MiD51 (Mitochondrial Dynamics Protein 51), MFF (Mitochondrial Fission Factor), and FIS1 (Fission, Mitochondrial 1), however, their distinct roles in fission are not completely understood (Losón et al., 2013; Osellame et al., 2016; Palmer et al., 2013). After the association with adaptor proteins, DRP1 then forms a multimeric ring around the mitochondria and hydrolyzes GTP to GDP causing a confirmational change in the ring that executes fission (Chang and Blackstone, 2007; Fröhlich et al., 2013; Nagashima et al., 2020). DRP1 is essential for mitochondrial fission and without functional DRP1, the mitochondrial network becomes hyperfused. In addition to the effects on mitochondrial morphology, DRP1 also mediates the fission of peroxisomes (Koch et al., 2003). Peroxisomes are highly specialized organelles vital for the beta oxidation of very-long-chain fatty acids, detoxification of reactive oxygen species (ROS), production of cholesterol, bile acids, and plasmalogens, which contribute to the phospholipid content in the brain white matter (Chandel, 2021; Ding et al., 2021; Ferreira et al., 2021; Lodhi and Semenkovich, 2014; Ravi et al., 2021; Schrader et al., 2020; Sugiura et al., 2017). The adaptor proteins MFF and FIS1 are also involved in DRP1 recruitment to sites of peroxisome fission (Gandre-Babbe and Van Der Bliek, 2008; Itoyama et al., 2013; Koch and Brocard, 2012). The contacts between mitochondria and peroxisomes are essential for linking peroxisomal β-oxidation and mitochondrial ATP generation by the electron transport chain (ETC), maintenance of redox homeostasis, and lipid transfer (Chandel, 2021; Fan et al., 2016; Fransen et al., 2017). Thus, peroxisomal and mitochondrial metabolic activities are functionally interconnected. Consistent with this, peroxisomal disorders caused by mutations in genes involved in the biogenesis and function of peroxisomes, are often associated with mitochondrial dysfunction (Tanaka et al., 2019). This communication may be particularly important during brain development and metabolic stress. However, the molecular mechanisms that facilitate this communication are still poorly understood.

In mouse models, global DRP1 knockout is embryonic lethal (Ishihara et al., 2009; Wakabayashi et al., 2009). With the advent of exome sequencing, patients with *de novo* mutations in the gene that encodes DRP1, *DNM1L (Dynamin 1 Like)*, have been identified (Chao et al., 2016; Fahrner et al., 2016; Keller et al., 2021; Liu et al., 2021; Longo et al., 2020; Nasca et al., 2016; Nolan et al., 2019; Ryan et al., 2018; Sheffer et al., 2016; Vandeleur et al., 2019; Waterham et al., 2007; Yoon et al., 2016). The disease associated with mutations in *DNM1L* is known as <u>e</u>ncephalopathy due to defective <u>m</u>itochondrial and <u>p</u>eroxisomal fission-1 (EMPF1) (OMIM 614388). Most reported mutations are heterozygous missense mutations. Patients present with heterogeneous symptoms including neurodevelopmental delay, seizures, muscle abnormalities, and ataxia. Patient fibroblasts show very elongated mitochondrial networks and elevated lactate levels, suggesting an impairment in oxidative phosphorylation (Liu et al., 2021; Whitley et al., 2018). Currently, there is no effective treatment or cure for patients effected by DRP1 mutations. The life expectancy for most patients with DRP1 mutations is childhood to early adolescence (Ryan et al., 2018). Although other proteins and organelles are involved in fission, DRP1 has been shown to be indispensable for mitochondrial fission (Csordás et al., 2006; Fonseca et al., 2019; Kamerkar et al., 2018; Li et al., 2015; Nagashima et al., 2020; Smirnova et al., 2001), yet the cellular consequences of DRP1 dysfunction are not well understood. In this study, we aimed to characterize the effects of two DRP1 mutations identified in patients with EMPF1, the G32A mutation, which lies in the catalytic GTPase domain of DRP1, and the R403C mutation, which lies in the stalk domain (Fröhlich et al., 2013) (Figure 1A), on mitochondrial network morphology, cristae ultrastructure, and related metabolic adaptations, to gain fundamental insight into the cellular adaptations to dysfunctional organelle fission and the pathophysiology of EMPF1.

**Figure 1.**
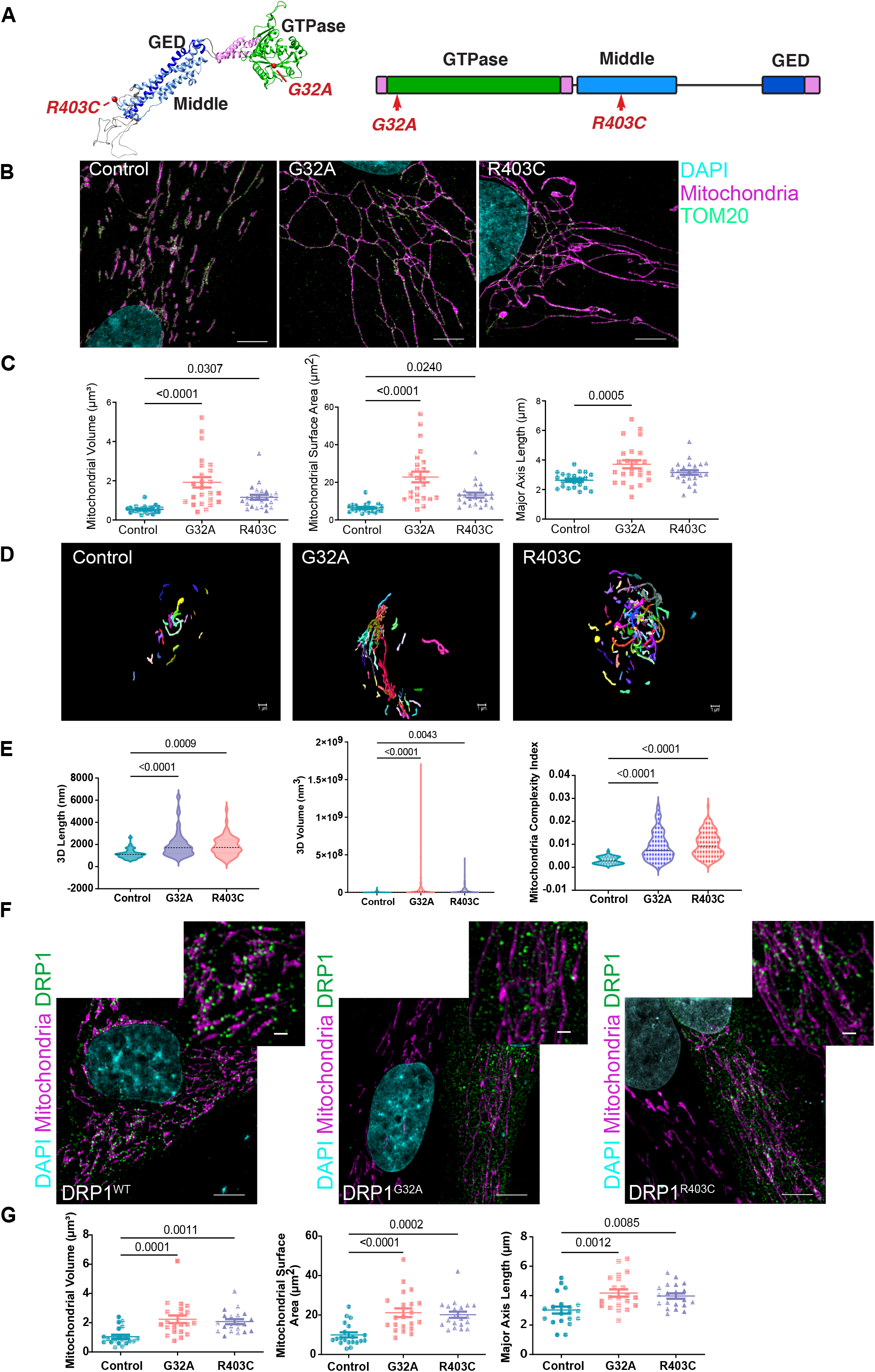
DRP1 patient-derived fibroblasts have elongated mitochondrial morphology. (A) Structural and schematic representation of DRP1 with mutations highlighted in red. (B) Representative images of mitochondrial morphology in patient-derived fibroblasts using structured illumination microscopy (n=3, 7 cells per replicate). (C) Quantification of mitochondrial volume, surface area, and major axis length. Analyzed using one-way ANOVA followed by Dunnett’s multiple comparison test. (D) Representative sbf-SEM 3D reconstructed images. Each mitochondrion is colored a unique color. (E) Quantification of sbf-SEM for 3D mitochondrial length, volume, and complexity index. Analyzed using one-way ANOVA followed by Dunnett’s multiple comparison test. (F) Representative images of mitochondrial morphology in control fibroblasts transfected with either DRP1^WT^-mCherry, DRP1^G32A^-mCherry, or DRP1^R403C^-mCherry using structured illumination microscopy (n=3, 7 cells per replicate). (G) Quantification of mitochondrial volume, surface area, and major axis length. Analyzed using one-way ANOVA followed by Dunnett’s multiple comparison test.

Here, we resolve and quantify mitochondrial and peroxisomal morphological features associated with EMPF1-associated mutations and reveal their impact on cellular metabolism, to put into perspective the clinical findings that were previously described (Fahrner et al., 2016; Liu et al., 2021; Ryan et al., 2018). We found that the inability of DRP1 to fragment the mitochondria in patient-derived fibroblasts leads not only to the hyperfusion phenotype previously described, but also to delayed cell death in response to apoptotic signals, aberrant mitochondrial cristae structure, and hyperpolarized mitochondrial membrane potential. Metabolically, the changes in mitochondrial and peroxisomal morphology result in inefficiency of ETC coupling oxygen consumption with ATP production and upregulation of glycolysis. Altogether, these data indicate that mitochondrial and peroxisomal fission influence dynamics of the inner mitochondrial membrane and the metabolic state of the cell.

## Results

### DRP1 variants found in patients have dominant negative effects that alter the expression of critical proteins at the mitochondria

Light microscopy studies have shown the profound alterations of mitochondrial fission in cells from patients harboring mutations in DRP1 (Ryan et al., 2018; Whitley et al., 2018). These studies highlighted the clinical relevance of mitochondrial fission in neural development. We quantified the morphology of the mitochondria in patient-derived primary fibroblasts at super resolution level using structured illumination microscopy (SIM). Both patient fibroblast lines showed increased mitochondrial volume, surface area and major axis length, consistent with a hyperfused mitochondrial morphology, as reported for DRP1 null cells (Fonseca et al., 2019; Kamerkar et al., 2018; Smirnova et al., 2001) (Fig. 1B-C). Live-cell imaging of mitochondria stained with MitoTracker confirmed that mitochondrial fission events are rare in patient cells compared to control fibroblasts (Supplementary Fig. 1A). High resolution three-dimensional images of control and patient fibroblasts were obtained using serial block facing-scanning electron microscopy (SBF-SEM) (Garza-Lopez et al., 2021). Manual segmentation and stack reconstruction of the mitochondrial networks validated the increased length and volume of the mitochondrial networks as well as increased mitochondrial complexity index (MCI) in patient fibroblasts (Fig. 1D-E). MCI is a quantitative measurement of network complexity, which takes into account both volume and surface area of the mitochondria (MCI = SA^3^ / 16π^2^V^2^) (Faitg et al., 2021; Vincent et al., 2019). Both mutations in DRP1 result in significant alterations of the mitochondrial network with G32A fibroblasts showing slightly more severe elongation than R403C (Fig. 1E and Supplementary Fig. 1B).

Both mutations in DRP1 are found to be heterozygous in patients, suggesting that mutant DRP1 has a dominant effect that can interfere with the ability of wild-type DRP1 to fragment the mitochondria. To test this, we generated mCherry tagged- constructs containing the patient derived mutations and expressed them in human fibroblasts homozygous for endogenous wild-type DRP1. Overexpression of mCherry-DRP1-G32A and mCherry-DRP1-R403C by transient transfection in wild-type fibroblasts demonstrated that, as previously described (Montecinos-Franjola et al., 2020, Fahrner et al., 2016; Ryan et al., 2018; Whitley et al., 2018), the mitochondrial hyperfusion seen in patient cells is not due to the genetic background and that these mutations act in a dominant negative manner (Fig. 1F-G). While both mutations of DRP1 significantly impaired DRP1 function in human fibroblasts, the G32A show a slightly more severe phenotype than R403C.

Hyperfusion of the mitochondrial membrane due to sequestration and inactivation of DRP1 is thought to lead to unopposed fusion events that increase the severity of mitochondrial elongation (Palmer et al., 2013). We examined the effects of DRP1 mutations on the protein expression levels of mitochondrial dynamics proteins. Both Mitofusin 1 and Mitofusin 2 were expressed at lower levels in patient cells than in control cells. However, there was no significant difference in the expression of OPA1 (Fig. 2A). These data show that there is a reduction in the expression of mitochondrial outer membrane fusion proteins in patient cells, suggesting that active mitochondrial fission promotes protein stability at the outer mitochondrial membrane and compensatory fusion events.

**Figure 2.**
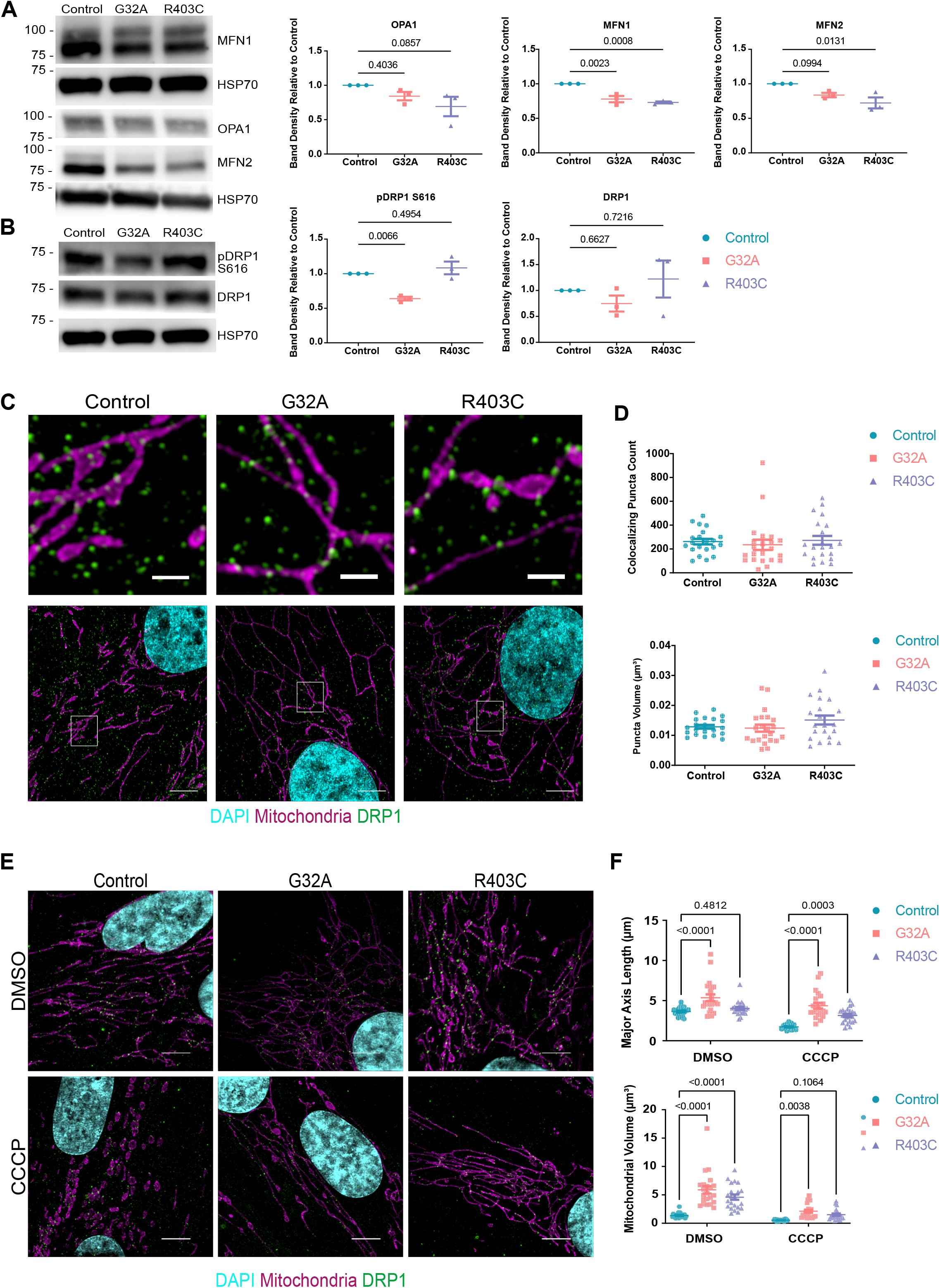
Mutant G32A and R403C DRP1 are recruited to mitochondria. (A) Western blot of total protein lysate isolated from patient fibroblasts and probed for mitochondrial fusion proteins. Representative image of three independent biological replicates. Band density is normalized to loading control (HSP70) and relative to control cells. (B) Western blot of total protein lysate isolated from patient fibroblasts and probed for mitochondrial fission proteins. Band density is normalized to loading control (HSP70) and relative to control cells. Representative image of three independent biological replicates. (C) Representative images of mitochondrial and DRP1 in patient-derived fibroblasts using structured illumination microscopy (n=3, 7 cells per replicate). Top image is a zoom of white box from bottom image. (D) Quantification of DRP1 puncta that overlap with mitochondria and volume of DRP1 puncta. Analyzed using one-way ANOVA followed by Dunnett’s multiple comparison test. (E) Representative images of mitochondrial morphology in patient-derived fibroblasts treated with either DMSO or CCCP for 1 hours using structured illumination microscopy (n=3, 7 cells per replicate). (F) Quantification of mitochondrial volume, surface area, and major axis length. Analyzed using one-way ANOVA followed by Dunnett’s multiple comparison test.

Phosphorylation of DRP1 at Serine 616 has been proposed to be a critical determinant of DRP1 recruitment and activity (Koch and Brocard, 2012; Taguchi et al., 2007; Valera-Alberni et al., 2021). Whether mutations in DRP1 found in patients with EMPF1 affect the levels of phosphorylation at Serine 616 has not been explored. We examined DRP1 levels in patient derived fibroblasts. Although there was no significant difference in total DRP1 expression, G32A patient cells expressed significantly less pS616-DRP1 than control cells (Fig. 2B). Since G32A mutation impairs DRP1 phosphorylation to a greater extent than the R403C mutation, the GTPase domain of DRP1 may be structurally or functionally involved in DRP1 phosphorylation. We wondered whether the expression of one copy of mutant DRP1 in patient fibroblasts affected the recruitment of endogenous pS616-DRP1 to the mitochondria. To test this, we quantified the recruitment of phosphorylated DRP1 to the outer mitochondrial membrane using super-resolution microscopy. On average, there was no significant difference in the recruitment of pS616-DRP1 to the mitochondria in patient cells and control cells (Fig. 2C-D). While in all cell lines, pS616-DRP1 was found in close contact to the mitochondrial membrane, in the patient fibroblasts this interaction appeared less transient. These results point to generalized stalling of the fission events in patient fibroblasts.

The onset of apoptosis is accompanied by mitochondrial remodeling, stable association of DRP1 to the mitochondrial membrane, and increased fragmentation of the mitochondrial membrane (Martinou and Youle, 2006; Wasiak et al., 2007). In DRP1 deficient cells, this remodeling of mitochondria is significantly reduced and the onset of cell death is delayed (Lee et al., 2004). To determine the effects of mitochondrial stress on mitochondrial fragmentation in patient fibroblasts, we quantified changes in mitochondrial fission in response to CCCP treatment. As previously shown using RNAi experiments in HeLa cells, patient fibroblasts showed reduced mitochondrial fragmentation induced by CCCP compared to control cells (Fig. 2E-F). Although some mitochondrial fragmentation may occur, mitochondrial volume and major axis length are significantly larger in patient cells treated with CCCP than control cells. Upon apoptosis, BAX translocates to the mitochondria and cytochrome *c* is released to the cytosol (Smaili et al., 2001). While in DRP1 depleted cells BAX is recruited to the mitochondria, cytochrome *c* release is delayed. Consistent with this, patient fibroblasts show higher cell survival when treated with CCCP and Etoposide; as well as, lower caspase-3 activity than control fibroblasts when treated with STS (Supplementary Fig. 2A-B). Interestingly, basal levels of both BAX and BAK are significantly reduced in patient fibroblasts with mutations in DRP1 (Supplementary Fig. 2C). Levels of the anti- apoptotic protein MCL-1 were significantly reduced in G32A fibroblasts but not in R403C fibroblasts, while BCL-XL expression levels remained relatively constant in all cell lines. Thus, patient fibroblasts may display a compensatory adaptation to reduce the basal levels of BAX/BAK activation. Whether this reduction in the ability to undergo apoptosis in response to mitochondrial stress is a widespread phenomenon in all tissues of EMPF1 patients and/or whether apoptosis inhibition results in deleterious accumulation of mitochondrial and nuclear DNA encoded mutations remains unexplored. No significant alteration in the protein expression of DRP1 adaptors was detected (Supplementary Fig. 2D).

### Mitochondrial hyperfusion in DRP1 mutant fibroblasts leads to higher glycolytic rate and increased mitochondrial membrane potential

Mitochondrial morphology is tightly linked to metabolic function (Dorn, 2015; Wai and Langer, 2016); thus, we assessed the effects of DRP1 patient mutations on cellular metabolism by measuring rates of glycolysis and oxidative phosphorylation (OXPHOS) in cells using the sensitive high-throughput Seahorse XF Analyzer. The Seahorse analyzer measures extracellular oxygen consumption as a readout of OXPHOS and extracellular acidification as a readout of glycolysis. Basal and maximal oxygen consumption was not significantly different in mutant cells compared to control, suggesting no major defects in ATP production (Fig. 3A). However, proton leak was significantly higher in the G32A patient cells and coupling efficiency was significantly lower in both patient lines (Fig. 3B). Therefore, although basal ATP production is not perturbed in the mutant cells, the ETC is not effectively coupling substrate oxidation and ATP synthesis. Extracellular acidification rate was significantly increased in both patient lines, consistent with an upregulation of glycolysis (Fig. 3A) and the hyperlactacidemia observed in more than 50% of patients with mutations in DRP1 (Liu et al., 2021). A glycolytic rate assay confirmed that both basal and compensatory glycolysis are higher in both patient lines (Fig. 3C). Thus, although the cells are capable of producing ATP, DRP1 mutations cause a defect in OXPHOS efficiency and a slight metabolic dependency on glycolysis (Fig. 3C and Supplementary Fig. 3A).

**Figure 3.**
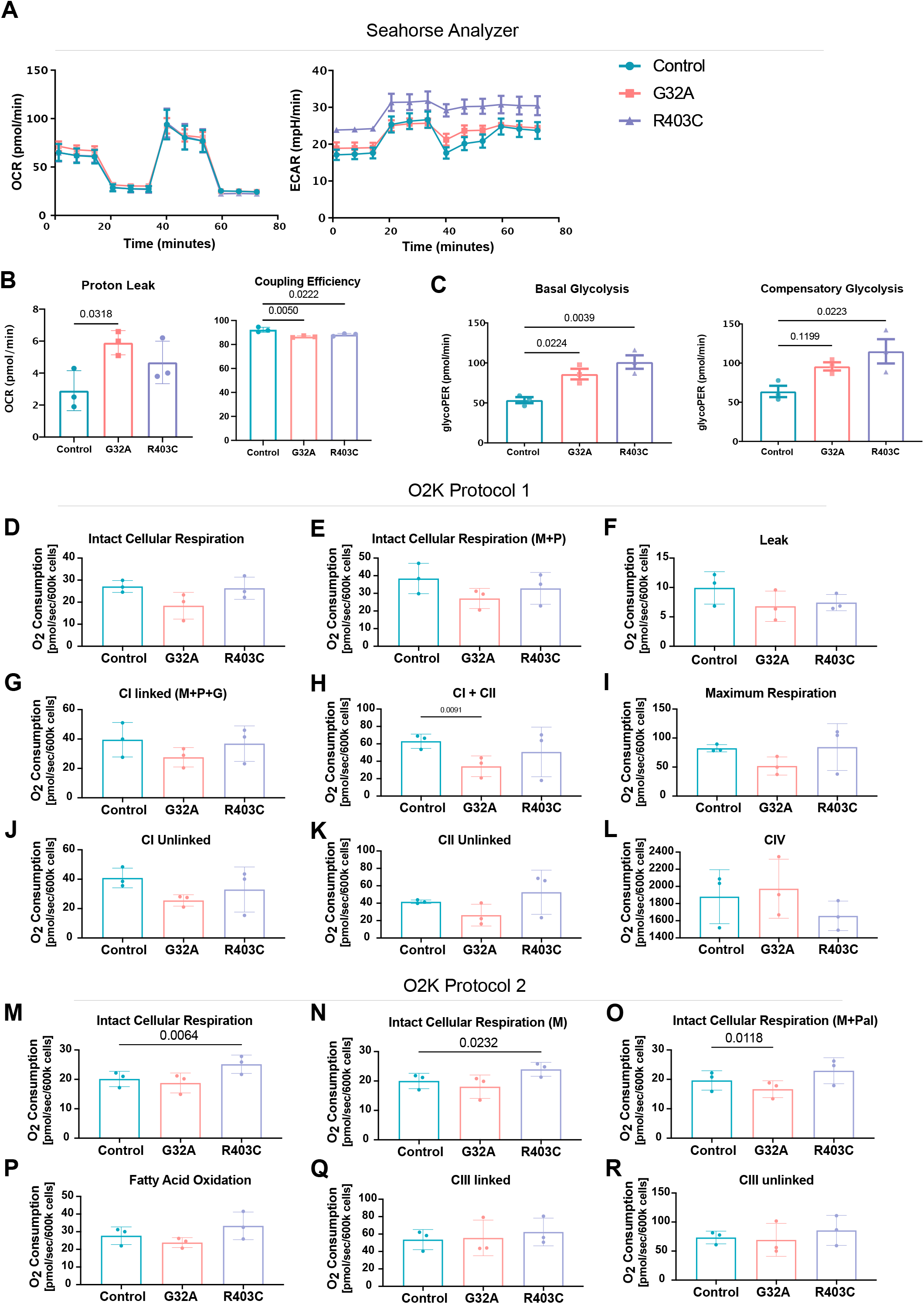
Mitochondrial defects lead to inefficient coupling of the electron transport chain and higher glycolytic rate. (A) Oxygen consumption rate and extracellular acidification rate in patient fibroblasts as measured using Seahorse Mitochondrial stress test (n=3, 20 wells per replicate). Oligomycin applied at 20 minutes, FCCP applied at 40 minutes, and rotenone/Antimycin A applied at 60 minutes. Analyzed using two-way ANOVA followed by Šídák’s multiple comparisons test. (B) Proton leak and coupling efficiency calculated using oxygen consumption rate. (C) Basal glycolysis of patient fibroblasts measure using Seahorse Glycolytic Rate Assay. Compensatory glycolysis measured following application of rotenone/Antimycin A. (D-L) O_2_ consumption rate of intact cells followed by isolated complex O_2_ consumption using protocol 1. Analyzed using one-way ANOVA followed by Dunnett’s multiple comparison test. (M-R) O_2_ consumption rate of intact cells followed by isolated complex O_2_ consumption using protocol 2. Analyzed using one-way ANOVA followed by Dunnett’s multiple comparison test.

To dissect functional differences in the ETC complexes at the mitochondria between control and mutant fibroblasts, we complemented our studies with the high-resolution Oxygraph-2k (O2k, Oroboros Instruments, Austria) which measures OXPHOS within intact, permeabilized cells (Ye and Hoppel, 2013). This method includes two protocols run in parallel (Hsiao and Hoppel, 2018; Krahenbuhl et al., 1991; Ye and Hoppel, 2013) using a number of selective substrates and inhibitors to measure the rates of oxygen consumption attributed to distinct ETC complexes. In protocol 1, the activity of complexes I, II, and IV is sequentially assessed and measured. First, intact cellular respiration (ICR) is determined to obtain a baseline rate prior to the addition of any substrates. Consistent with the Seahorse analyzer, the DRP1 mutant fibroblasts showed equivalent baseline OXPHOS to control fibroblasts (Fig. 3D). Then, substrates for Complex I (CI), malate (M) and pyruvate (P), were added, but elicit no impact on the cellular respiration as these substrates remain extracellular (Fig. 3E). To permeabilize the cell membrane, digitonin (Dig) was added, resulting in diluted intracellular content and allowing M + P to enter the cell for oxidation, which reveals the leak rate in the absence of ADP. In this assay, there was not a significant increase in proton leak between control and mutant fibroblasts when substrate was not limited (Fig. 3F). Based on our results from Seahorse where the major phenotype between the mutant lines and the control fibroblasts was in the decreased coupling efficiency of the mitochondria, we examined the functionality of each of the ETC complexes. Thus, adenosine diphosphate (ADP) was added, to stimulate respiration of coupled mitochondria (also known as state 3)(Hsiao and Hoppel, 2018; Ye and Hoppel, 2013), followed by addition of glutamate (G), another CI substrate, and succinate (S), a complex II substrate. The addition of these compounds provides an assessment of the coupled CI OXPHOS and the coupled CI + CII OXPHOS. While we found no significant differences in the CI capacity (Fig. 3G), the DRP1 G32A fibroblasts showed a significant decrease in the coupled CI/CII OXPHOS (Fig. 3H). We then assess the maximal oxidative capacity, by adding the uncoupler, trifluoromethoxy carbonylcyanide phenylhydrazone (FCCP), and saw a slight decrease, albeit not significant in the DRP1 G32A fibroblasts (Fig. 3I). Next, the addition of rotenone (Rot), an inhibitor of CI, assesses the uncoupled CI and CII rates individually. No significant differences were detected (Fig. 3J-K). To assess complex CIV Oxphos, we added tetramethyl-p-phenylenediamine (TMPD) and ascorbate (AS), which donates electrons to cytochrome c (cyt c), then a CIV inhibitor, azide, was added. No differences in CIV activity were detected (Fig. 3L). Thus, our assessment of complexes I, II, and IV in the mutant G32A fibroblasts revealed a potential deficiency in the linked CI/CII mitochondrial OXPHOS. This decrease in linked CI/CII activity is likely due to the additive effect of combined rate since both CI and CII trended down on their own. This decreased in CI and CII activity was not seen in the R403C fibroblasts.

In protocol 2, we measured the OXPHOS for FAO and complex III (CIII). The ICR rate was first assessed (Fig 3M). Followed by the addition of Malate (M) and palmitoylcarnitine (Pal), a long-chain fatty acid derivative (Fig. 3N-O). We found that the ICR was significantly higher in the DRP1 R403C mutant fibroblasts (Fig. 3M-N) and significantly decreased in DRP1 G32A fibroblasts upon addition of M and P (Fig. 3O). Upon permeabilization with Digitonin, ADP was then added to assess the state 3 rate of Pal oxidation. No significant overall differences in the rate of FAO were detected under these conditions (Fig. 3P). We followed by the addition of Rot to diminish any non-mitochondrial oxidation, and duroquinol (DHQ), a CIII substrate, to assess the coupled CIII rate (Fig. 3Q). Finally, the uncoupled CIII rate was evaluated via the addition of FCCP. There were no differences in the coupled CIII activity or the uncoupled CIII oxidation. Collectively, our results showed impaired coupled CI/CII activity in G32A but not R403C fibroblasts, and a potential increase in FAO capacity in R403C mutants.

The upregulation of glycolysis with no accompanied increase in OXPHOS led us to interrogate a potential impairment in pyruvate transport. Two proteins are essential for mitochondrial pyruvate transport in humans, Mitochondrial Pyruvate Carrier 1 and 2 (MPC1 and MPC2) (Bricker et al., 2012; Herzig et al., 2012). MPC1 and MPC2 associate to form an ∼150-kilodalton complex in the inner mitochondrial membrane. Real time quantitative PCR (RT-qPCR) gene expression analysis showed a significant downregulation of *MPC2* expression in R403C cells and *MPC1* in both patient lines (Supplementary Fig. 3B). These data suggest that the upregulation in glycolysis seen in DRP1 mutant patient fibroblasts is accompanied by impaired pyruvate import into the mitochondria. We examined the expression of MPC1 and MPC2, using Western blot, which revealed a significant decrease in MPC2 expression in the R403C cells (Supplementary Fig. 3C)

The major contributor of the proton motive force of the ETC is the mitochondrial membrane potential, which drives OXPHOS. Thus, changes in coupling efficiency can point to perturbation of the mitochondrial membrane potential and the maintenance of key ion gradients. We investigated this possibility using the fluorescent sensor Tetramethylrhodamine, ethyl ester (TMRE). TMRE is a cell-permeant, cationic, red-orange fluorescent dye that is sequestered by polarized mitochondria. We found that mitochondrial membrane potential as reported by TMRE was significantly increased in patient fibroblasts (Fig. 4), indicative of hyperpolarization of the mitochondria.

**Figure 4.**
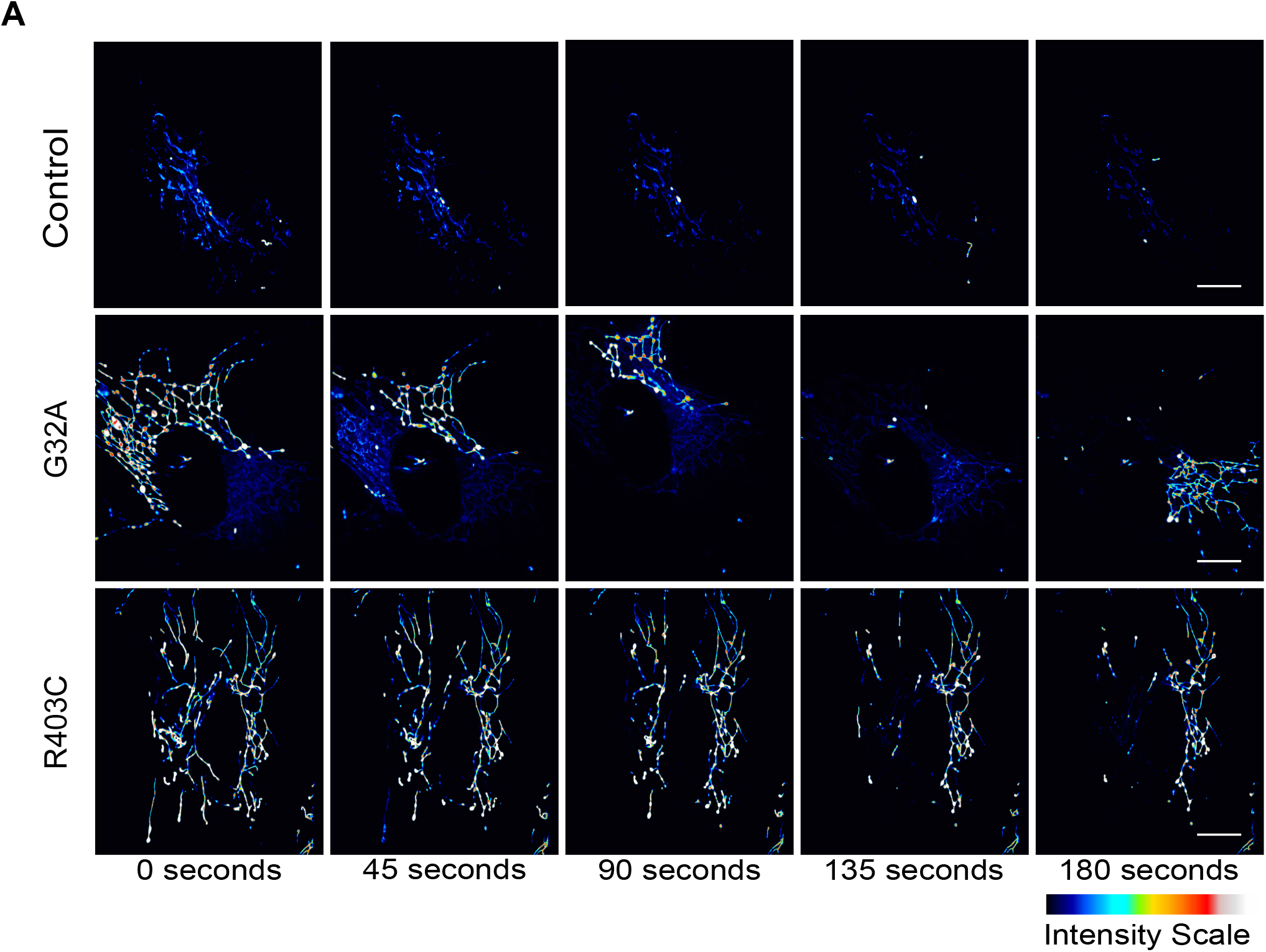
DRP1 patient fibroblasts have more hyperpolarized mitochondrial membrane potential. (A) Representative live-cell imaging of patient-derived fibroblasts treated with 500nM TMRE for 10 minutes. Cells were imaged every 15 seconds for 15 minutes using spinning disk confocal (n=3, 7 cells per replicate). Scale bar: 5 µm.

### DRP1 mutations alter mitochondrial cristae morphology

Cristae topology readjustments participate in energy balance control (Eramo et al., 2020; Kondadi et al., 2020). The hyperpolarization of the mitochondrial membrane detected in patient fibroblasts lead us to assess the effects of DRP1 mutations on cristae morphology. We used a previously optimized method to quantify and analyze cristae morphology changes from transmission electron microscopy (TEM) (Eisner et al., 2017; Lam et al., 2021). In this method, cristae abundance is calculated by manually quantifying the number of cristae per mitochondrion, and a score between 0 (worst) and 4 (best) is assigned based on the morphology: (0—no well-defined cristae, 1—no cristae in more than ∼75% of the mitochondrial area, 2—no cristae in approximately ∼25% of mitochondrial area, 3—many cristae (over 75% of area) but irregular, 4—many regular cristae) (Supplementary Fig. 4A). TEM of patient fibroblasts confirmed the mitochondrial elongation observed in SIM (Supplementary Fig. 4B-E) and revealed differences in the cristae structure (Fig. 5A). The quantification of cristae morphology demonstrated a reduced cristae score and decreased cristae number relative to mitochondrial length in DRP1-deficient cells compared with the control cells (Fig. 5B). These data link aberrant outer mitochondrial membrane fission caused by DRP1 mutations to generalized cristae structural changes. The structure and respiratory function of mitochondrial cristae are maintained by the <u>mi</u>tochondrial contact sites and <u>c</u>ristae <u>o</u>rganizing <u>s</u>ystem (MICOS) – a large complex of scaffolding and membrane shaping proteins localized at the inner mitochondrial membrane. Given the reduced cristae score, we examined expression of some key components of the MICOS complex. *MIC19, MIC25,* and *MIC60* are all reduced in either one or both patient lines at the mRNA level; however, protein expression is not significantly changed (Supplementary Fig. 4F). Thus, the structural abnormalities seen in patient fibroblasts are not due to changes in the expression of MICOS proteins. Additional biochemical assays are needed to assess the functionality of these proteins in the context of EMPF1.

**Figure 5.**
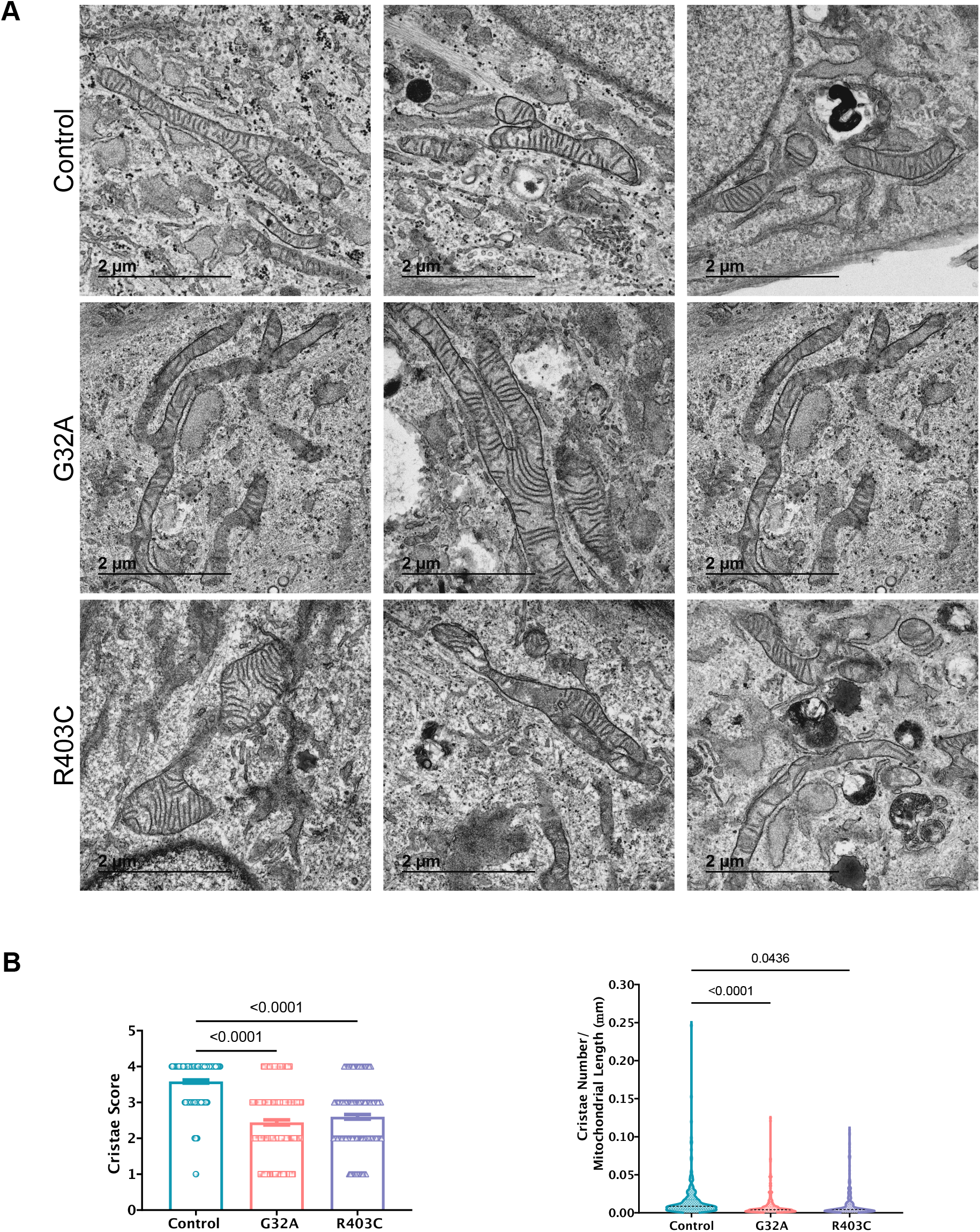
Cristae morphology is altered in patient cells. (A) Transmission electron microscopy of patient fibroblasts. 3 representative images per line shown. (B) Quantification of cristae from TEM images for cristae score and cristae number relative to mitochondrial length. Analyzed using one-way ANOVA followed by Dunnett’s multiple comparison test.

### Patient fibroblasts display hyperfused peroxisomes

In addition to catalyzing mitochondrial fission, DRP1 also participates in peroxisomal fission (Kamerkar et al., 2018). While fission of peroxisomes has been demonstrated to be involved in *de novo* peroxisomal biogenesis, the functional consequences of peroxisomal fission are not completely understood. Previous reports using EMPF1 patient fibroblasts have not reported elongation of peroxisomes, but DRP1 null cells have been reported to have hyperfused peroxisomes (Kamerkar et al., 2018). To interrogate the effects of DRP1 mutations on peroxisome morphology, we performed super-resolution microscopy. SIM revealed that both mutations in DRP1 result in significant elongation of peroxisomes (Fig. 6A). Quantification of the peroxisome structure shows an increase in volume, surface area, and length of peroxisomes (Fig. 6B), demonstrating that DRP1 mutations found in patients with EMPF1 result in impaired peroxisome fission.

**Figure 6.**
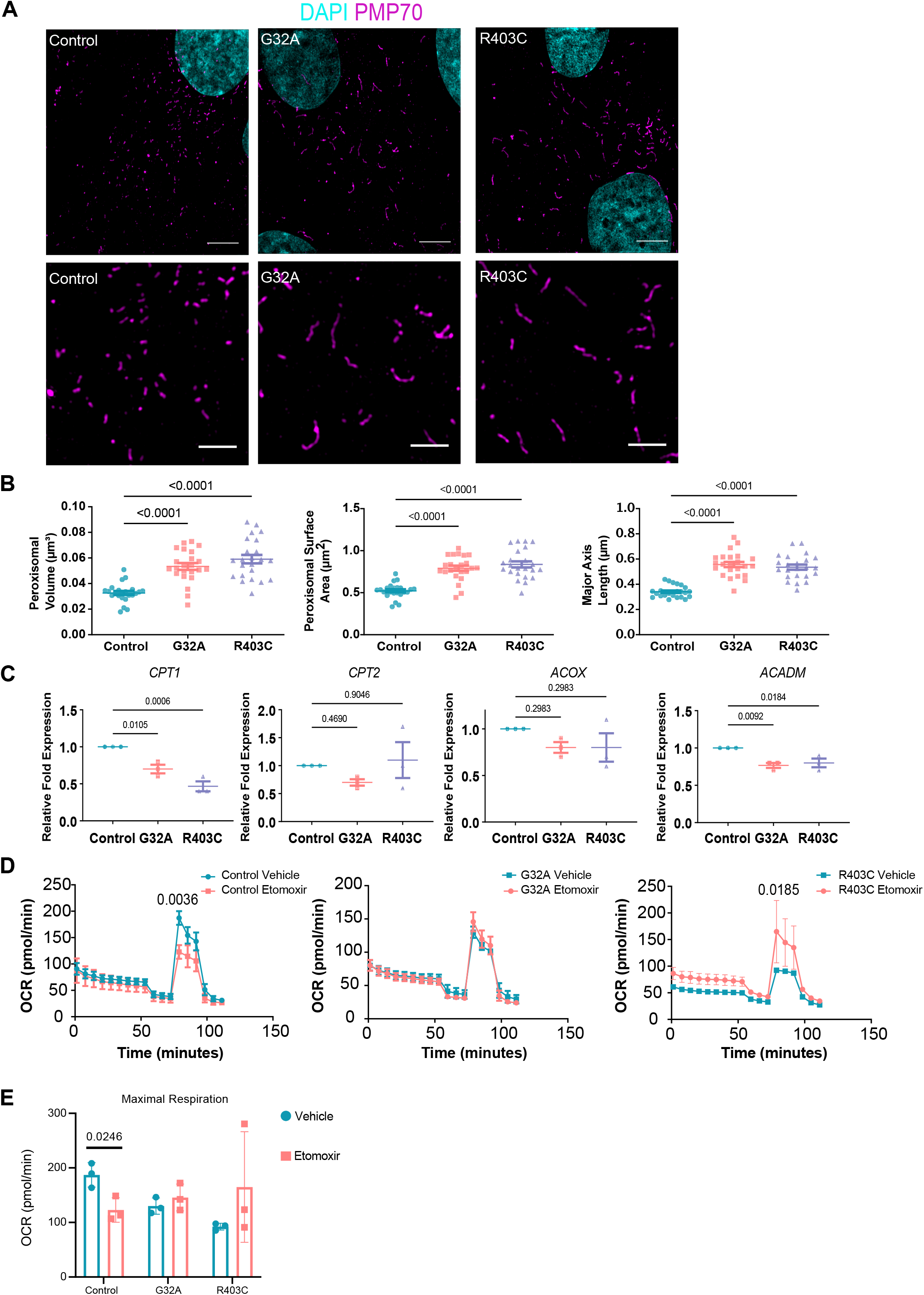
Patient fibroblasts have elongated peroxisomes. (A) Representative images of peroxisomal morphology in patient-derived fibroblasts using structured illumination microscopy (n=3, 7 cells per replicate). (B) Quantification of peroxisomal volume, surface area, and major axis length. Analyzed using one-way ANOVA followed by Dunnett’s multiple comparison test. (C) qRT-PCR analysis of gene expression in fibroblasts relative to control and normalized to two housekeeping genes (*GPI* and *GAPDH)*. Quantifications are shown for three independent biological replicates. (D) Oxygen consumption rate in patient fibroblasts as measured using Seahorse Analyzer XFe96 (n=3, 20 wells per replicate). Etomoxir applied after 20 minutes, oligomycin applied at 40 minutes, FCCP applied at 60 minutes, and rotenone/Antimycin A applied at 80 minutes. Analyzed using two-way ANOVA followed by Šídák’s multiple comparisons test. (E) OCR at maximal respiration after FCCP treatment.

Since peroxisomes and mitochondria are hubs for the regulation of fatty acid metabolism, we first tested whether the dramatic changes in morphology would alter the expression of key genes involved in fatty acid oxidation in patient fibroblasts. The expression of these genes have previously been reported to increase in a DRP1 phosphomimetic mouse model (Valera-Alberni et al., 2021). RT-qPCR analysis showed that the expression of carnitine palmitoyltransferase I, *CPT1*, and acyl-coA dehydrogenase medium chain, *ACADM*, are significantly lower in patient cells compared to control (Fig. 6C). CPT1 is associated with the outer mitochondrial membrane and is responsible for catalyzing the transfer of the acyl group of acyl-CoA to L-carnitine. Fatty acids are then able to move from the cytosol into the inner membrane space of the mitochondria and the matrix, where fatty acid oxidation occurs. *ACADM* encodes for the medium-chain-acyl-CoA dehydrogenase (MCAD) protein, which catalyzes the first step of mitochondrial fatty acid beta-oxidation. Thus, the reduction in *CPT1* and *ACADM* expression may hinder the ability of the mitochondria to carryout fatty acid oxidation.

To examine the ability of patient fibroblasts to perform fatty acid oxidation, we used the Seahorse analyzer to quantify oxygen consumption rates of patient fibroblasts treated with either vehicle or etomoxir, a CPT1 inhibitor. The cells were cultured with minimal glucose and glutamine and high L-carnitine and palmitate to promote fatty acid oxidation over glycolysis or glutaminolysis. Control fibroblasts treated with etomoxir, an inhibitor of CPT1, show a reduction in maximal respiration following FCCP injection. In contrast, the G32A patient-derived cells show no change in OCR with etomoxir treatment and the R403C patient-derived cells show an increase in maximal respiration following FCCP injection (Fig. 6D-E). These data indicate that in contrast to the reliance of control cells to long chain fatty acid oxidation under conditions of maximal respiration, both patient lines have lost this dependency and adapted to the inhibition of long-chain fatty acid import into mitochondria under these conditions. These data show that fibroblasts with DRP1 mutations appear to have compensatory adaptations to alterations in long-chain fatty acid oxidation.

### Mitochondrial hyperfusion is rescued by MFN2 silencing

The exact mechanisms by which the mitochondrial fusion machinery is positioned to sites of fusion are not completely understood. However, fission and fusion events occur in rapid succession (Liu et al., 2009) and fission and fusion machineries localize in close proximity at the endoplasmic reticulum membrane contact sites (Abrisch et al., 2020). Given the intricate relationship between these opposing dynamics, we examined the rates of fission and fusion in live cells to determine if defects in fission may impact fusion rates. Our analysis in live cells showed that in control cells the number of fission and fusion events per frame are very similar, supporting the concept that these dynamic events are in equilibrium and constantly occurring in tandem to one another under physiological conditions. By contrast, in patient cells, the reduced number of fission events was accompanied by a significantly decreased number of fusion events (Fig. 7A). Thus, defective fission leads to fewer fusion events contributing to decreased dynamic mitochondrial network phenotype seen in patient cells.

**Figure 7:**
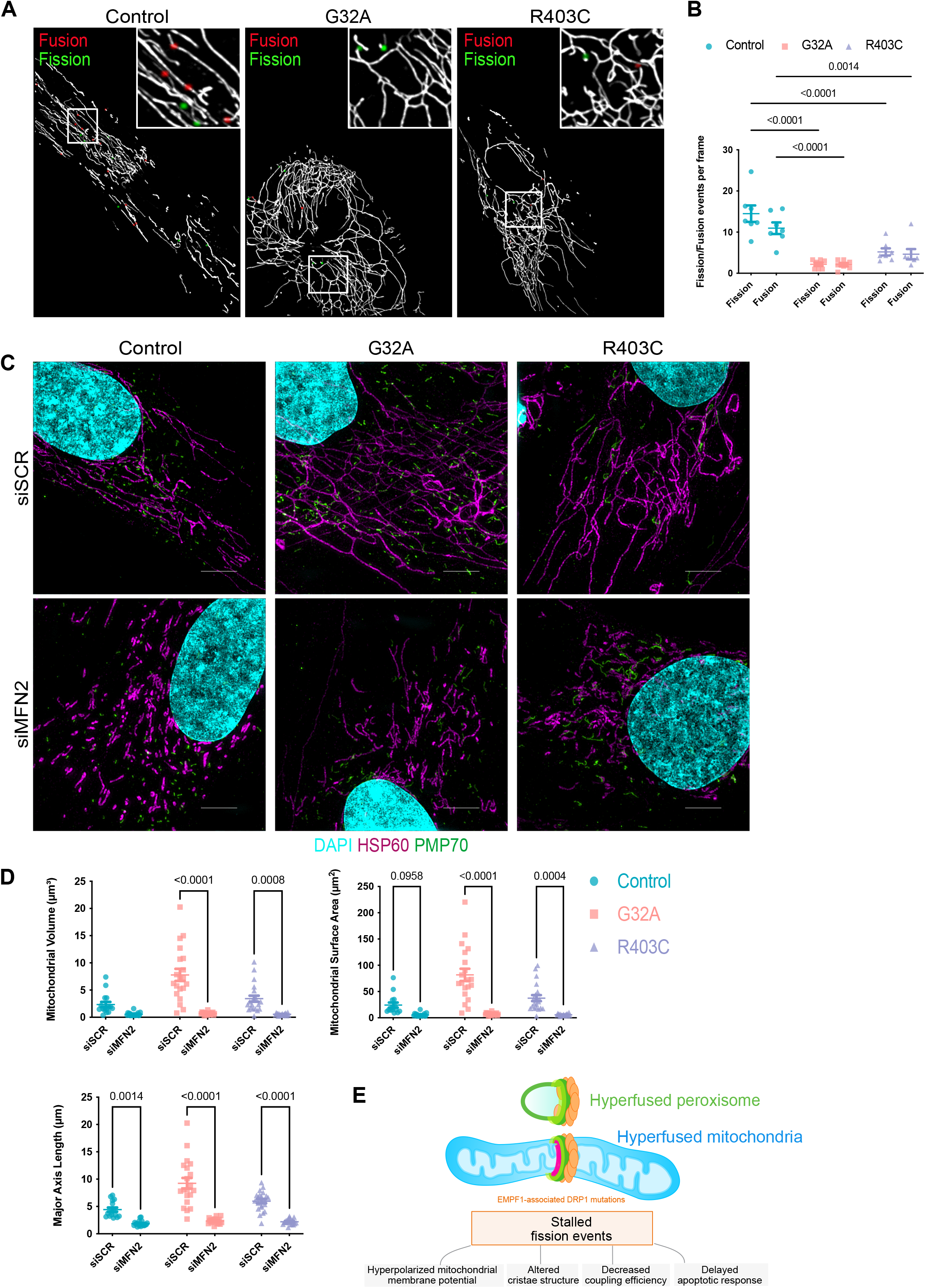
Current model of the cellular responses to EMPF1 associated DRP1 mutations. (A) Representative images of mitochondrial dynamics in patient-derived fibroblasts treated with Mitotracker Green with fission events marked in green and fusion events marked in red (7 cells per replicate). (B) Quantification of mitochondrial dynamics using MEL. Analyzed using one-way ANOVA followed by Dunnett’s multiple comparison test. (C) Representative images of mitochondrial and peroxisomal morphology in patient-derived fibroblasts treated with either siRNA against MFN2 or scrambled using structured illumination microscopy (n=3, 7 cells per replicate). (D) Quantification of mitochondrial volume, surface area, and major axis length. Analyzed using one-way ANOVA followed by Dunnett’s multiple comparison test. (E) Summary schematic: EMPF1- associated mutations do not appear to alter the capacity of DRP1 to be recruited to points of fission, but rather cause stalling of the fission events. Pausing peroxisomal and mitochondrial fission results in alterations of cristae structure, changes in mitochondrial membrane potential, decreased mitochondrial coupling efficiency, delayed cell death response, and overall metabolic adaptations.

The deleterious effects of fission deficiency have been shown to be restored by the concomitant deletion of the mitochondrial fusion machinery (Chen H., et al., 2015). To further determine the contribution of fusion to the mitochondrial network phenotype, we next tested whether deletion of MFN2 would rescue the mitochondrial hyperfusion phenotype in patient fibroblasts. Indeed, MFN2 deletion restored all the parameters of mitochondrial morphology altered by DRP1 mutations (Fig. 7B), while the peroxisomal elongation was not completely restored (Supplementary Fig. 5). Collectively these results show that the mitochondrial hyperfusion phenotype can be rescued by modulating the levels of mitochondrial fusion.

## Discussion

DRP1 is an essential gene for embryonic development (Ishihara et al., 2009; Wakabayashi et al., 2009). RNAi based approaches in human cell culture models have been used to explore the impact of DRP1 loss on mitochondrial fission and downstream processes such as apoptosis and mitophagy (Kleele et al., 2021; Otera et al., 2016; Smirnova et al., 2001). Although loss of DRP1 results in mitochondrial hyperfusion, it’s not fully understood how mutations in DRP1 can impact cellular metabolic function, cristae structure, and/or peroxisomal function. The analysis of organelle morphology in fibroblast lines derived from patients with EMPF1 that harbor one wild-type copy of DRP1 and one missense copy of DRP1, demonstrate a dominant negative effect of mutant DRP1 that results in stalling of the fission machinery that has a wide impact on cellular function.

Mitochondrial fission is essential for cellular processes such as apoptosis, mitophagy, and mitochondrial biogenesis. However, how mitochondrial fission helps maintain cristae structure and/or cellular metabolism have not been thoroughly explored. Our studies revealed that while DRP1 is recruited to the mitochondria in EMPF1 patient cell lines, the fission event is incomplete, and thus, mitochondrial fission is paused. This stalling of mitochondrial fission results in dramatic changes to cristae morphology, notably a decrease in cristae score and number, indicative of immature and poorly formed cristae. Changes in redox signaling can lead to cysteine oxidation of MIC60 (Li et al., 2021; Wasilewski and Chacinska, 2021), an inner mitochondrial membrane protein responsible for cristae structure integrity. Thus, the lack of proper mitochondrial fission in patient cells may disrupt redox signaling, which may in turn be responsible for the aberrant cristae morphology detected in patient cells.

Detailed metabolic analysis of patient fibroblasts showed that these cells have decreased coupling efficiency and upregulation of glycolysis, suggesting that the functional impact of stalled fission is electron transport chain inefficiency. This inefficiency may be due to the changes in cristae structure, since cristae structure is essential for F_0_F_1_–ATPase to function effectively (Afzal et al., 2021; Cogliati et al., 2013; Davies et al., 2011; Wilkens et al., 2013). Previous studies revealed that there are distinct fission signatures that are key determinants of the decision of mitochondria to proliferate or degrade (Kleele et al., 2021). Peripheral fission events were shown to be preceded by a decrease in membrane potential and proton motive force. We found that the elongated mitochondria from patient fibroblasts have a hyperpolarized membrane potential, which could result from inefficient function of the F_0_F_1_–ATPase. Under certain conditions, the F_0_F_1_– ATPase can pump protons out of the mitochondrial matrix into the intermembrane space in a reverse-mode action, causing ΔΨm hyperpolarization. Alternatively, hyperpolarization may be due to complex I inhibition or VDAC (voltage-dependent anion channel) opening, which can cause hyperpolarization independently of F_0_F_1_–ATPase reverse-mode action (Forkink et al., 2014). A comprehensive analysis of the functionality of the ETC complexes would be needed to elucidate the connection between fission stalling events and mitochondrial hyperpolarization. Upregulation of glycolysis in these patient fibroblasts could result in subsequent glucose depletion locally within the cells, which could in turn cause ΔΨm hyperpolarization. Carbon-tracing could help reveal whether all of the products of glycolysis are efficiently shuttled into the TCA cycle or if intermediates are lost in the process. Investigating the function of uncoupling proteins such as UCP2 and UCP3 may elucidate whether the decreased efficiency is a byproduct or cause of mitochondrial dysfunction, such as increased ΔΨm, in these cells. Our finding that cells with DRP1 mutations have profound alterations in cristae morphology but apparent maintenance of overall ATP production, raises the interesting possibility that DRP1 mutant cells have evolved specialized mechanisms to cope with electron leakage and damaging reactive oxygen species. A landmark study from Spinelli and collaborators demonstrated the ability of complex II to work in reverse in some tissues with the intrinsic capacity to use fumarate as a terminal electron acceptor (Spinelli et al., 2021). It is plausible that complex II reversal has evolved as a mechanism to maintain mitochondrial function in cells with alterations in fission caused by DRP1 mutations.

The existence of two mechanistically and functionally distinct types of fission at the mitochondria and the peroxisomes have not been clearly elucidated. In our study, we demonstrate that patient fibroblasts with mutations in DRP1 also have elongated peroxisomal morphology. Previous studies in mammalian cell culture revealed that peroxisome membrane elongation precedes membrane constriction and final membrane fission (Delille et al., 2010; Schrader et al., 2016). Whether peroxisomes have a specialized fusion machinery is still under debate. However, we speculate that under particular metabolic conditions, peroxisomes, particularly those in contact with mitochondria, may be able to fuse to respond to cellular metabolic needs. Elongation of the peroxisomes in various human disorders has been associated with alterations in fatty acid oxidation (Waterham et al., 2016). However, in these cells, peroxisomal function cannot be fully dissociated from changes in mitochondrial morphology. Our results suggest that MFN2 silencing could serve as a useful tool to discern the contribution of peroxisomal elongation independently of the mitochondrial network. Our model (Fig. 7) suggests that the mechanistic and functional connection between peroxisome remodeling and metabolic homeostasis could be dysregulated in mitochondrial diseases such EMPF1. We propose that in this context, human diseases could provide insight into the causal mechanisms underlying basic cellular processes, such as organelle fission. The detailed elucidation of DRP1-centric mechanisms will reveal the connection between organelle dynamics and cell fate. Several questions remain: Does stalling of mitochondrial fission affect the association/dissociation of other proteins at the site of fission? Is peripheral fission also inhibited in patient fibroblasts? If that is the case, what are the consequences for mitophagy and mitochondrial DNA integrity? What are the effects of peroxisomal elongation on fatty acid metabolism? How is cell division and mitochondrial segregation affected by DRP1 mutations? How are cell fate transitions affected by DRP1 mutations? Answering these fundamental questions would provide insight into the mechanisms of mitochondrial fission under the lens of this rare disease caused by dysregulated mitochondrial and peroxisomal homeostasis and inform the field on more rational and specific therapeutic targets for these devastating neurodevelopmental disorders.

## STAR Methods

### Cell Culture

Human fibroblasts were thawed and plated directly in maintenance medium consisting of MEM (Gibco #11095098) with 10% FBS (Sigma-Aldrich #F2442) and non-essential amino acids (Gibco # 11-140-050). Cells were maintained at 37°C and 5% CO2 with media changes every other day. Cells were passaged as needed using TrypLE (Thermo Fisher #12-064-013). For imaging experiments, cells were re-plated onto #1.5 glass-bottom 35 mm dishes (Cellvis #D35C4-20-1.5- N).

### Cell Treatments

All treatments were added directly to fibroblasts in maintenance medium. The electron transport chain decoupler CCCP (Sigma-Aldrich #C2759-250MG) was added to cells at a concentration of 1 µM. Cells were treated for 2 hours prior to fixation and imaging. All stock solutions were prepared in DMSO.

### Plasmid Transfection

Plasmid encoding mCherry-DRP1 (Addgene #49152) was transfected using FuGENE (Promega #E2311) as described in the manufacturer protocol. Patient mutations were introduced to plasmids using the In Fusion cloning kit (Takara Bio #638909) as described in the manufacturer protocol. Cells were maintained until optimal transfection efficiency was reached before cells were imaged.

### siRNA Transfection

siRNA knockdown against MFN2 (Thermo Fisher #s19260) or negative control scrambled was performed using Lipofectamine RNAiMAX tranfection reagent (Thermo Fisher # 13778150). Cells were fixed and stained 48hrs after transfection.

### Immunofluorescence

For immunofluorescence, cells were fixed with 4% paraformaldehyde for 20 min and permeabilized in 1% Triton-X-100 for 10 min at room temperature. After blocking in 10% BSA, cells were treated with primary and secondary antibodies using standard methods. Cells were mounted in Fluoromount-G (Electron Microscopy Sciences #17984-25) prior to imaging. Primary antibodies used include mouse anti-Mitochondria (Abcam #ab92824), rabbit anti-Tom20 (Cell Signaling Technology #42406S), rabbit anti-DRP1 (Cell Signaling Technology #8570S), rabbit anti-HSP60 (Cell Signaling Technologies # 12165S), mouse anti-PMP70 (Thermo Fisher # MA5- 31368), and rabbit anti-pDRP1 S616 (Cell Signaling Technology #3455S). Secondary antibodies used include donkey anti-mouse Alexa Fluor-488 (Thermo Fisher Scientific #A21202), donkey anti-rabbit Alexa Fluor-488 (Thermo Fisher Scientific #A21206), and donkey anti-mouse Alexa Fluor-546 (Thermo Fisher Scientific #A10036), and donkey anti-rabbit Alexa Fluor-546 (Thermo Fisher Scientific #A10040).

### Western Blot

Gel samples were prepared by mixing cell lysates with LDS sample buffer (Life Technologies, #NP0007) and 2-Mercaptoethanol (BioRad #1610710) and boiled at 95°C for 5 minutes. Samples were resolved on 4-20% Mini-PROTEAN TGX precast gels (BioRad #4561096) and transferred onto PVDF membrane (BioRad #1620177). Antibodies used for Western blotting are as follows: rabbit anti-DRP-1 (Cell Signaling Technologies #8570S), rabbit anti-pDRP-1 S616 (Cell Signaling Technologies #3455S), rabbit anti-OPA1 (Cell Signaling Technologies #67589S), mouse anti- MFN1 (Abcam #ab57602, mouse anti-MFN2 (Abcam #ab56889), rabbit anti-MFF (Proteintech #17090-1-AP), rabbit anti-MiD49 (Proteintech #16413-1-AP), rabbit anti-MiD51 (Proteintech #20164-1-AP), and rabbit anti-FIS1 (Proteintech #10956-1-AP), and mouse anti-HSP70 (Invitrogen #MA3-006).

### Seahorse XF Analysis

Fibroblasts were plated onto Seahorse XF96 V3 PS cell culture microplates 2 days before the assay at 50,000 cells per well. One hour prior to the assay, media was switched to XF DMEM media containing 1 mM pyruvate, 2 mM glutamine, and 10 mM glucose. For the Seahorse XF Mito Stress Test, oxygen consumption rate (OCR) was measured sequentially after addition of 1.0 μM oligomycin, 0.5 μM FCCP, and 0.5 μM rotenone plus Antimycin A. Protein leak and coupling efficiency were calculated using OCR. For the Seahorse XF Glycolytic Rate Assay, basal glycolytic rate was measured by extracellular acidification, and compensatory glycolysis was measured after addition of 0.5 μM rotenone plus antimycin A. Treatment with 50 mM 2-DG was used as a negative control. For the Seahorse XF Fuel Flex Test, glutamine dependency was measured by sequential addition of 3.0 μM BPTES followed by 4.0 μM Etomoxir plus 2.0 μM UK5099. Fatty acid dependency was measured by sequential addition of 4.0 μM Etomoxir followed by 3.0 μM BPTES plus 2.0 μM UK5099. Glucose dependency was measured by sequential addition of 2.0 μM UK5099 followed by 3.0 μM BPTES plus 4.0 μM Etomoxir. For the Seahorse XF Long Chain Fatty Acid Oxidation Stress Test, 24 hours prior to assay cells were switched to base media containing 0.5 mM glucose, 1.0 mM glutamine, 0.5 mM XF L-carnitine, and 1.0% fetal bovine serum. One hour prior to the assay, media was switched to XF DMEM media containing 2 mM glucose, 0.5 mM L-Carnitine, and Palmitate-BSA. Oxygen consumption rate (OCR) was measured sequentially after addition of 4.0 μM etomoxir, 1.5 μM oligomycin, 2.0 μM FCCP, and 0.5 μM rotenone plus Antimycin A.

### O2K Analysis

Oxygen consumption was measured with an O2K system (OROBOROS) using Protocols 1 and 2 described in Ye and Hoppel, 2013. Briefly, 2 ml of cell suspension (600,000 cells/ml) in Mir05 buffer was added to each chamber. Initial substrates were added (malate and pyruvate in protocol 1 and malate and palmitoylcarnitine in protocol 2) followed by addition of digitonin in increments (2ug/ml) to permeabilize the cell membranes. The final concentration of substrates and inhibitors was composed of malate (2 mM), pyruvate (2.5 mM), adenosine diphosphate (ADP, 2.5 mM), glutamate (10 mM), succinate (10 mM), palmitoylcarnitine (10 μM), duroquinol (0.5 mM), etramethyl-p-phenylenediamine (TMPD, 0.5 μM), ascorbate (2 mM), carbonylcyanide-p-trifluoromethoxy- phenylhydrazone (FCCP, 0.5-μM increment), rotenone (75 nM), antimycin A (125 nM), and sodium azide (200 mM). For Protocol 1, the non-mitochondrial (AA-insensitive) rate was subtracted. For Protocol 2, the Rot-insensitive rate was subtracted. For DHQ, the AA was subtracted.

### RNA Extraction and cDNA Synthesis

Cells were washed with PBS and scraped into 1000μl Trizol reagent. 200μl of chloroform was added and the samples were incubated at room temperature for 5 minutes. The samples were spun down at 12,000 g and the aqueous phase was collected. 500μl of isopropanol was added to precipitate RNA and the sample was incubated for 25 minutes at room temperature, followed by centrifugation at 12,000 g for 10 min at 4°C. The RNA pellet was washed with 75% ethanol, semi-dried, and resuspended in 30μl of DEPC water. After quantification and adjusting the volume of all the samples to 1μg/μl, the samples were treated with DNAse (New England Biolabs, # M0303). 10μl of this volume was used to generate cDNA using the manufacturer’s protocol (Thermo Fisher, #4368814).

### Quantitative RT PCR (RT-qPCR)

1ug of cDNA sample was used to run RT-qPCR for the primers mentioned in the table. QuantStudio 3 Real-Time PCR machine, SYBR green master mix (Thermo Fisher, #4364346), and manufacturer instructions were used to set up the assay.

### Image Acquisition

Super-resolution images were acquired using a Nikon SIM microscope equipped with a 1.49 NA 100x Oil objective an Andor DU-897 EMCCD camera. Live-imaging experiments were acquired on a Nikon Eclipse Ti-E spinning disk confocal microscope equipped with a 1.40 NA 60X Oil objective and an Andor DU-897 EMCCD camera. Image processing and quantification was performed using Fiji.

### Sample Preparation for Transmission Electron Microscope (TEM) Analysis

Human fibroblasts were immediately washed in ice-cold saline. Samples were processed as described by (Lam et. al., 2021, Pereira et al., 2017). Cells were fixed in 2.5% glutaraldehyde in sodium cacodylate buffer for 1 hour at 37 °C, embedded in 2% agarose, postfixed in buffered 1% osmium tetroxide, stained in 2% uranyl acetate, and dehydrated with an ethanol graded series. After embedding in EMbed-812 resin, 80 nm sections were cut on an ultramicrotome and stained with 2% uranyl acetate and lead citrate. Images were acquired on a JEOL JEM-1230 Transmission Electron Microscope, operating at 120 kV. Random images of human fibroblasts were obtained.

### TEM Analysis

Measurements of mitochondrial area, circularity, and length were performed using the multi-measure region of interest (ROI) tool in ImageJ (Lam et. al., 2021, Parra et al., 2014). To measure cristae morphology, we used ROIs in ImageJ to determine the cristae area, circulatory index, number, volume, and cristae score. The NIH ImageJ software was used to manually trace and analyze all mitochondria or cristae using the freehand told in the ImageJ application (Lam et. al., 2021, Parra et al., 2014). To assign a mitochondrion a cristae score, cristae abundance and form were evaluated. A score between 0 and 4 (0 – no sharply defined crista, 1 – greater than 50% of mitochondrial area without cristae, 2 – greater than 25% of mitochondrial area without cristae, 3 – many cristae but irregular, 4 – many regular cristae) was assigned (Eisner et al., 2017).

### Serial Block Facing-Scanning Electron Microscopy (SBF-SEM) Processing

Human fibroblasts were fixed in 2% glutaraldehyde in 0.1 M cacodylate buffer and processed using a heavy metal protocol adapted from a previously published protocol (Courson et al., 2021; Mustafi et al., 2014). Human fibroblasts were immersed in 3% potassium ferrocyanide and 2% osmium tetroxide (1 hour at 4 °C), followed by filtered 0.1% thiocarbohydrazide (20 min), 2% osmium tetroxide (30 min), and left overnight in 1% uranyl acetate at 4 °C (several de-ionized H2O washes were performed between each step). The next day samples were immersed in a 0.6% lead aspartate solution (30 min at 60 °C and dehydrated in graded acetone (as described for TEM). Human fibroblasts were impregnated with epoxy Taab 812 hard resin, embedded in fresh resin, and polymerized at 60 °C for 36–48 hours. After polymerization, the block was sectioned for TEM to identify the area of interest, then trimmed to 0.5 mm × 0.5 mm and glued to an aluminum pin. The pin was placed into an FEI/Thermo Scientific Volumescope 2 scanning electron microscope (SEM). The scope is a state-of-the-art SBF imaging system. For 3D EM reconstruction, thin (0.09 µm) serial sections, 300 to 400 per block, were obtained from the blocks that were processed for conventional TEM. Serial sections were collected onto formvar coated slot grids (Pella, Redding CA), stained, and imaged, as described above. The segmentation of SBF-SEM reconstructions was performed by manually tracing structural features through sequential slices of the micrograph block. Images were collected from 30–50 serial sections, which were then stacked, aligned, and visualized using Amira to make videos and quantify volumetric structures.

### Mitochondrial Event Localizer (MEL)

The MEL algorithm processes a fluorescence microscopy time-lapse sequence of z-stack images of Mitotracker- labeled mitochondria and produces event counts per frame and 3D locations indicating where mitochondrial events are likely to have occurred at each time step(Theart et al., 2020). These locations can subsequently be superimposed on the z-stacks to indicate the different mitochondrial events. The algorithm has been organized into two consecutive steps, namely the image pre-processing step which normalizes and prepares the time-lapse frames, and the automatic image analysis step which calculates the location of the mitochondrial events based on the normalized frames. MEL also allows each event to be subsequently validated and removed from the visualization and event counts. The code is available for download: https://github.com/rensutheart/MEL-Fiji-Plugin.

### Statistical Analysis

All experiments were performed with a minimum of 3 biological replicates. Statistical significance was determined by one-way or repeated measures one-way analysis of variance (ANOVA) as appropriate for each experiment. significance was assessed using Fisher’s protected least significance difference test. GraphPad PRISM v8.1.2and Statplus software package were used GraphPad Prism was used for all statistical analysis (SAS Institute: Cary, NC, USA). and data visualization. Error bars in all bar graphs represent standard error of the mean unless otherwise noted in the figure. No outliers were removed from the analyses. Quantification of mitochondrial morphology was performed in NIS-Elements (Nikon); briefly, we segmented mitochondria in 3D and performed skeletonization of the resulting 3D mask. Skeleton major axis length, volume, and surface area measurements were exported into Excel. For all statistical analyses, a significant difference was accepted when P < 0.05.

## Acknowledgements

We would like to thank the patients and families for permission to publish this work. We thank Dr. Eric Payne and Dr. Suzanne Hoppins for providing the patient-derived fibroblast lines used in this paper. We would also like to thank Dr. Evan Krystofiak for providing EM expertise, and Dr. Matthew Tyska, Dr. Uri Manor, and Dr. Kari Seedle, for providing advice and expertise with high-resolution microscopy. This work was supported by 1R35 GM128915-01NIGMS (VG), 1RF1MH123971-01 (VG); the Precision Medicine and Mental Health Initiative sponsored by the Vanderbilt Brain Institute (VG), 1R21CA227483-01A1NCI (VG), F99NS125829 (GLR), HHMI Gilliam (GLR), and CBMS T32 (GLR). All SIM and spinning disk confocal microscopy imaging and image analysis were performed in part through the use of the Vanderbilt Cell Imaging Shared Resource, which is supported by NIH grants 1S10OD012324-01 and 1S10OD021630-01. The authors declare no competing financial interests.

## Author Contributions

G. Robertson and V. Gama conceived the study, designed experiments, interpreted data, and wrote the manuscript. V. Gama supervised the research. G. Robertson designed and carried out all the cell biology experiments, with technical assistance from M. Patel and S. Riffle. J. Mears and M. Stoll performed the O2K experiments and provided technical feedback and insight into the conception and execution of study. A. Marshall, H. Beasley, Z. Vue, J. Shao, and E. Garza Lopez performed the TEM, 3D EM and sbf-SEM experiments, quantification, and analysis, with the supervision of A. Hinton. S. de Wet, R. P. Theart, and Ben Loos, applied the MEL algorithm to the mitochondrial imaging data to calculate fission and fusion events. All authors edited the document.

