## Supplemental Figure Legends for "DRP1-mediated mitochondrial fission is essential to maintain cristae morphology and bioenergetics"

**Supplementary Figure Legends**

**Supplementary Figure 1: DRP1 patient fibroblasts rarely undergo mitochondrial fission events.** (A) Live-imaging of mitochondria in patient fibroblasts using MitoTracker (n=3, 5 cells per replicate). Cells were imaged every 15 seconds for a total of 30 minutes. Scale bar: 1 µm. (B) Quantification of mitochondrial perimeter and 3D area from sbf-SEM.

**Supplementary Figure 2: DRP1 patient fibroblasts express equal levels of adaptor proteins but downregulate pro-apoptotic proteins.** (A) Cell titer blue assay with results relative to vehicle treated cells of same genotype. Analyzed using two-way ANOVA with Dunnett's multiple comparison test. (B) Caspase-Glo 3/7 assay with results relative to vehicle treated cells of same genotype. Analyzed using two-way ANOVA with Dunnett's multiple comparison test. (C) Western blot of total protein lysate isolated from patient fibroblasts and probed for pro and anti-apoptotic proteins. (D) Western blot of total protein lysate isolated from patient fibroblasts and probed for mitochondrial fission adaptor proteins. All quantified relative control and normalized to loading control (HSP70).

**Supplementary Figure 3: DRP1 patient fibroblasts rely more on glucose for fuel and downregulate expression of the mitochondrial pyruvate carriers.** (A) Glucose, glutamine, and fatty acid dependency of the electron transport chain measured using Seahorse Fuel Flex Assay. Glucose dependency measured after application of UK5099, glutamine dependency measured after application of BPTES, and fatty acid dependency measure after application of Etomoxir. (B) qRT-PCR analysis of gene expression in fibroblasts relative to control and normalized to two housekeeping genes (*GPI* and *GAPDH)*. Quantifications are shown for three independent biological replicates. (C) Western blot of total protein lysate isolated from patient fibroblasts and probed for mitochondrial pyruvate carriers. Band density is normalized to loading control (HSP70) and relative to control cells. Representative image of three independent biological replicates.

**Supplementary Figure 4: TEM cristae scoring examples and gene expression of MICOS complex components.** (A) Representative TEM images of mitochondria with cristae fitting the given cristae score. (B) Quantification of TEM images for mitochondria number, (C) mitochondrial length, (D) mitochondrial area, and (E) mitochondrial circularity index. (F) qRT-PCR analysis of gene expression in fibroblasts relative to control and normalized to two housekeeping genes (*GPI* and *GAPDH*).

**Supplementary Figure 5: Peroxisomal morphology is mostly unchanged after MFN2 knockdown.** Quantification of peroxisomal volume, surface area, and major axis length. Analyzed using one-way ANOVA followed by Dunnett's multiple comparison test.
