## Supplementary figures and images for "DRP1-mediated mitochondrial fission is essential to maintain cristae morphology and bioenergetics"

### Supplemental Figure 1

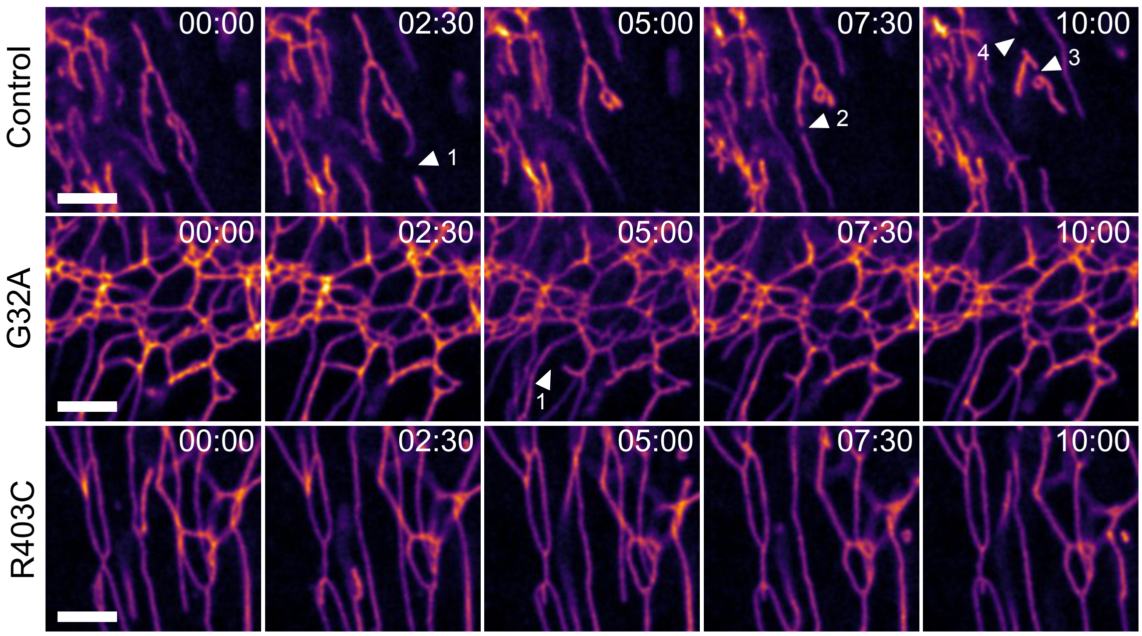

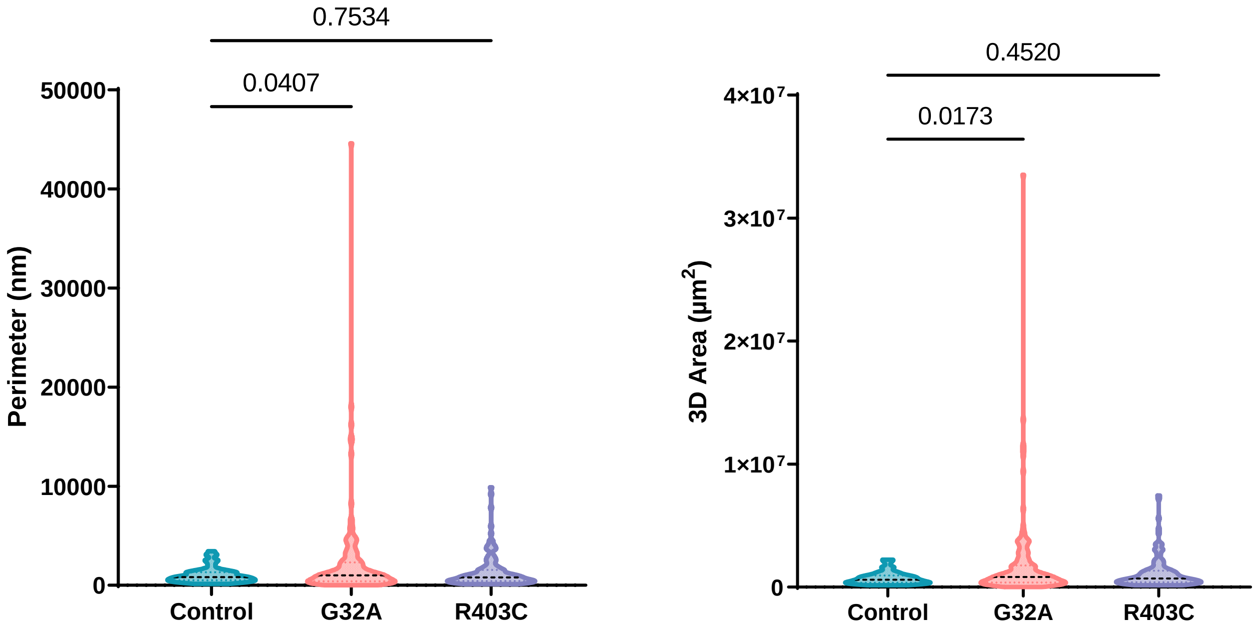

### Supplemental Figure 2

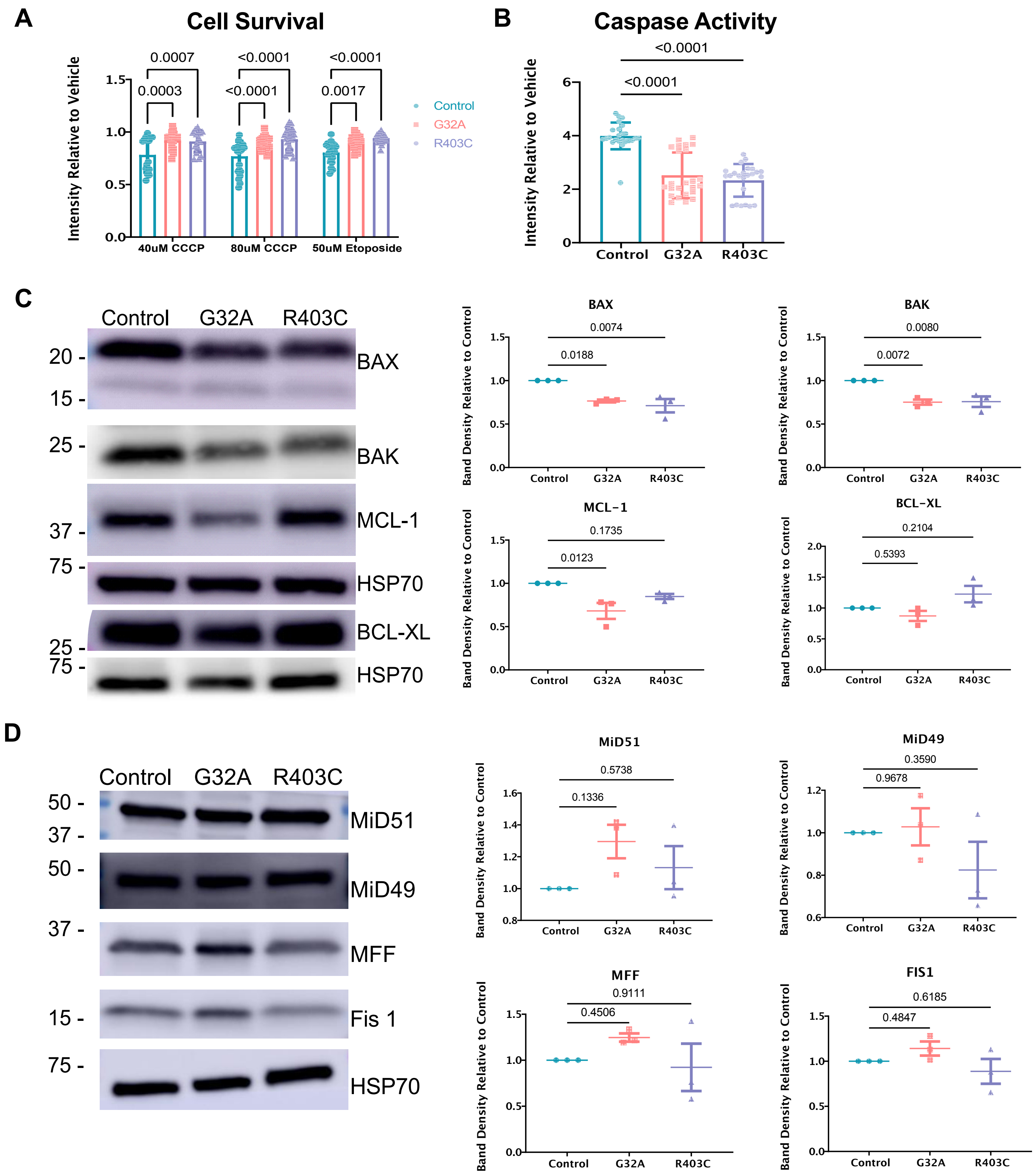

### Supplemental Figure 3

Supplementary Figure 3

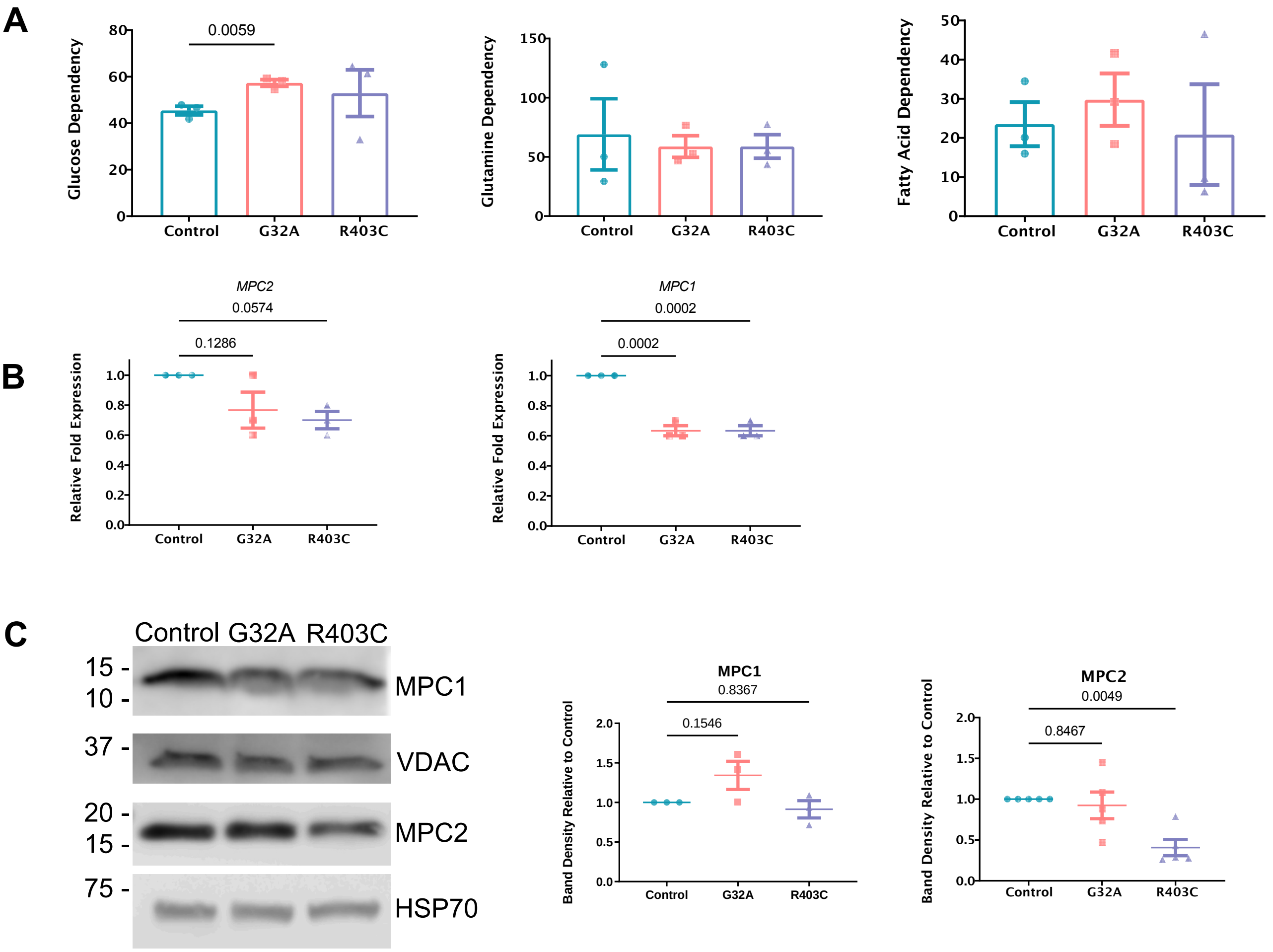

### Supplemental Figure 4

Supplementary Figure 4

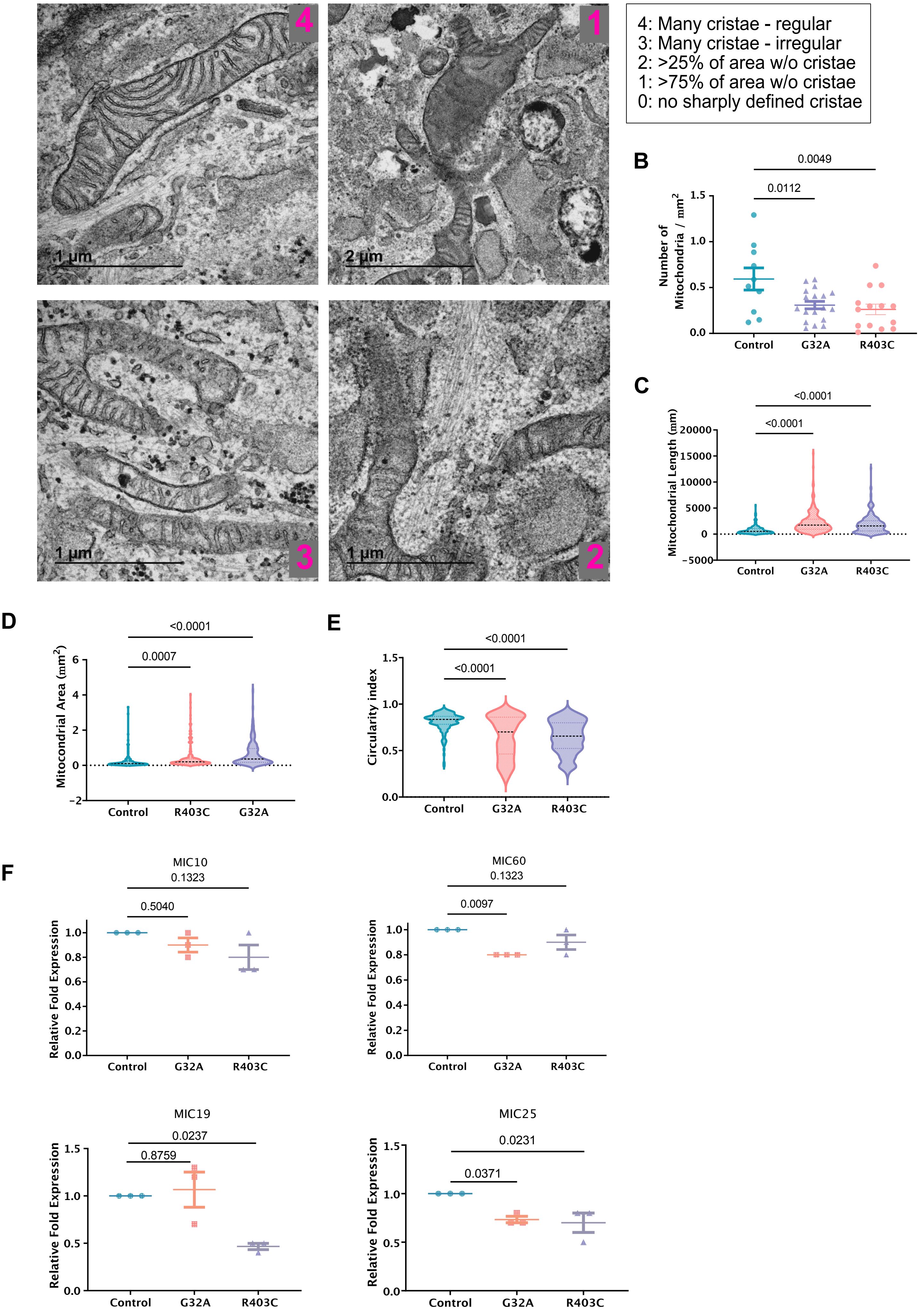

### Supplemental Figure 5

Supplementary Figure 5

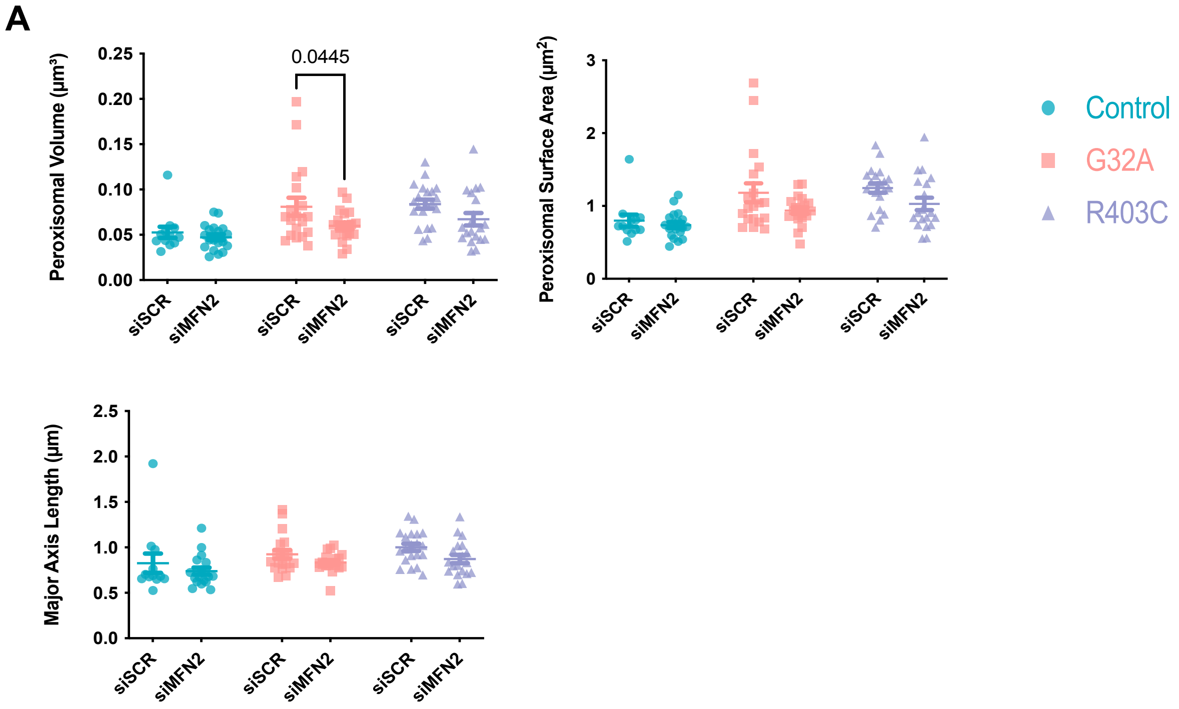
